# BioVix: An Integrated Large Language Model Framework for Data Visualization, Graph Interpretation, and Literature-Aware Scientific Validation

**DOI:** 10.64898/2026.02.03.703467

**Authors:** Muhammad Zain Butt, Rana Sheraz Ahmad, Eman Fatima, Muhammad Tahir ul Qamar

## Abstract

The application of Large Language Models (LLMs) for generating data visualizations through natural language interaction represents a promising advance in AI-assisted scientific analysis. However, existing LLM-based tools largely emphasize graph generation, while research workflows require not only visualization but also rigorous interpretation and validation against established scholarly evidence. Despite advances in visualization technologies, no single tool currently integrates literature references with visualization while also generating insights from graphical data. To address this gap, we present BioVix, a web-based LLM-driven framework that integrates interactive data visualization, natural-language querying, and automated retrieval of relevant academic literature. BioVix enables users to upload datasets, generate complex visualizations, interpret graphical patterns, and contextualize findings through literature references within a unified workflow. The system employs a multi-model architecture combining DeepSeek V3.1 for code and logic generation, Qwen2.5-VL-32B-Instruct for multimodal interpretation, and GPT-OSS-20B for conversational reasoning, coordinated through structured prompt engineering. BioVix was evaluated across diverse biological domains, including proteomic expression profiling, epigenomic peak annotation, and clinical diabetes data, demonstrating its flexibility in handling heterogeneous datasets and supporting exploratory, literature-aware analysis. While BioVix substantially streamlines exploratory research workflows, its LLM-generated outputs are intended to support, not replace, expert judgment, and users should independently verify results before scientific reporting. BioVix is openly available via public deployment on Hugging Face (https://huggingface.co/spaces/MuhammadZain10/BioVix), with source code provided through GitHub (https://github.com/MuhammadZain-Butt/BioVix).

## 1. Introduction

There are different methods for presenting data insights to a general audience, such as descriptive, predictive, and visual representations. However, the most user-friendly way to understand diverse patterns or trends is through interactive visualization (IV) [1]. This method allows people from diverse disciplines, including business, healthcare, social sciences, and education, to present existing trends, patterns, and insights derived from their data [1, 2]. With these generated IVs, even non-technical individuals can understand complex insights from data without requiring technical knowledge. Visualization methods have evolved through different stages. Conventionally, IVs relied heavily on programming expertise, typically in R or Python, along with specialized data visualization packages such as Matplotlib [3], Plotly [4], and ggplot2 [5]. This technical dependency has been a significant limitation for users without a programming background, often resulting in slower analysis and difficulty in understanding data trends. To address this drawback, an alternative and more accessible approach has emerged through the use of Natural Language Interfaces (NLIs) for data visualization [6]. NLIs enable users to create IVs by specifying Natural Language Queries (NLQs) rather than relying on programming. This approach has increased access to data visualization tools and has helped non-technical users to visualize and interpret data trends more independently [7].

The development of NLI-based visualization tools [8] has progressed through three primary stages, advancing language understanding and user-data interaction [9]. The initial stage was characterized by reliance on heuristics, rules-based approaches, and probabilistic grammar checking. These early approaches were constructed using rigid frameworks that used pre-defined grammatical structures and specialized vocabularies to handle user queries [10]. Tools, including [11], Deepeye [12], and FlowSense [13] employ these approaches during the early phase of NLI-based architectures to translate natural language (NL) input into IVs [14]. While these tools represented a significant advancement in NL-based IVs, their reliance on pre-defined linguistic rules imposed limitations that hindered the accurate satisfaction of user requirements. This often led to inaccurate visualizations, misinterpretation of user intent, and an inability to handle more complex queries [15]. The second stage of NLI-based visualization architectures marked a significant shift from rule-based approaches to deep neural network (DNN)-based approaches [16]. These deep learning models unified NL understanding, semantic reasoning, and visualization generation, enabling more flexible and robust NLI-based architectures [4]. These architectures were not based on fixed grammar rules or pre-defined instructions, unlike the previous approaches. Instead, these systems were trained on large datasets to acquire more detailed language conventions and semantic mappings between NLQs and the visual outputs. ADVISor [17], ncNet [18], and RGVisNet [19] are prominent tools that demonstrated the effectiveness of learning-based approaches in addressing the rigidity of linguistic rule-based tools [4]. The second stage enhanced the NLI-based visualization tools in handling user context, synonyms, and linguistic variations, which are essential for accurately addressing user queries [7]. Despite these advancements, progress remained constrained by the availability of large training datasets, limited model interpretability, and high computational requirements. These challenges pose substantial barriers to scalability and transparency. The current, third stage involves Large Language Models (LLMs), which have transformed the capabilities of NLI-based visualization tools [20]. These LLMs are built upon transformer frameworks that employ self-attention mechanisms to process complex user queries [21]. Chat2VIS represents one of the first LLM-based visualization tools that uses this paradigm, generating executable code to create IVs [14]. This approach marked a significant improvement in usability, allowing users to interact with data through a conversational interface [22]. Initially, these architectures were primarily driven by visualization tasks and conversion of NL syntax into specific code for creating IVs [23]. Currently, no single tool connects insights derived from generated visualizations with academic literature to validate observed data trends. As a result, users must manually search the literature to investigate trends or patterns identified in their graphs. This process is time-consuming, requiring users to first interpret the visualization, then formulate a research query, and finally retrieve relevant literature [9].

To address this, we developed BioVix, an LLM-based web tool that aims to integrate data visualization with academic literature **(Figure 1)**. BioVix allows users to generate IVs, obtain AI-assisted interpretations, formulate research queries, and retrieve the most relevant literature articles to analyze data trends observed in visualizations. In addition, BioVix employs state-of-the-art LLMs specialized for specific tasks, including DeepSeek V3.1, GPT-OSS-20B, and Qwen2.5-VL-32B-Instruct, for the visualization-to-literature search workflow, data-driven conversational support, and interpretation of uploaded graphs, respectively. We evaluated BioVix on three different datasets to demonstrate its efficiency and applicability.

**Figure 1.**
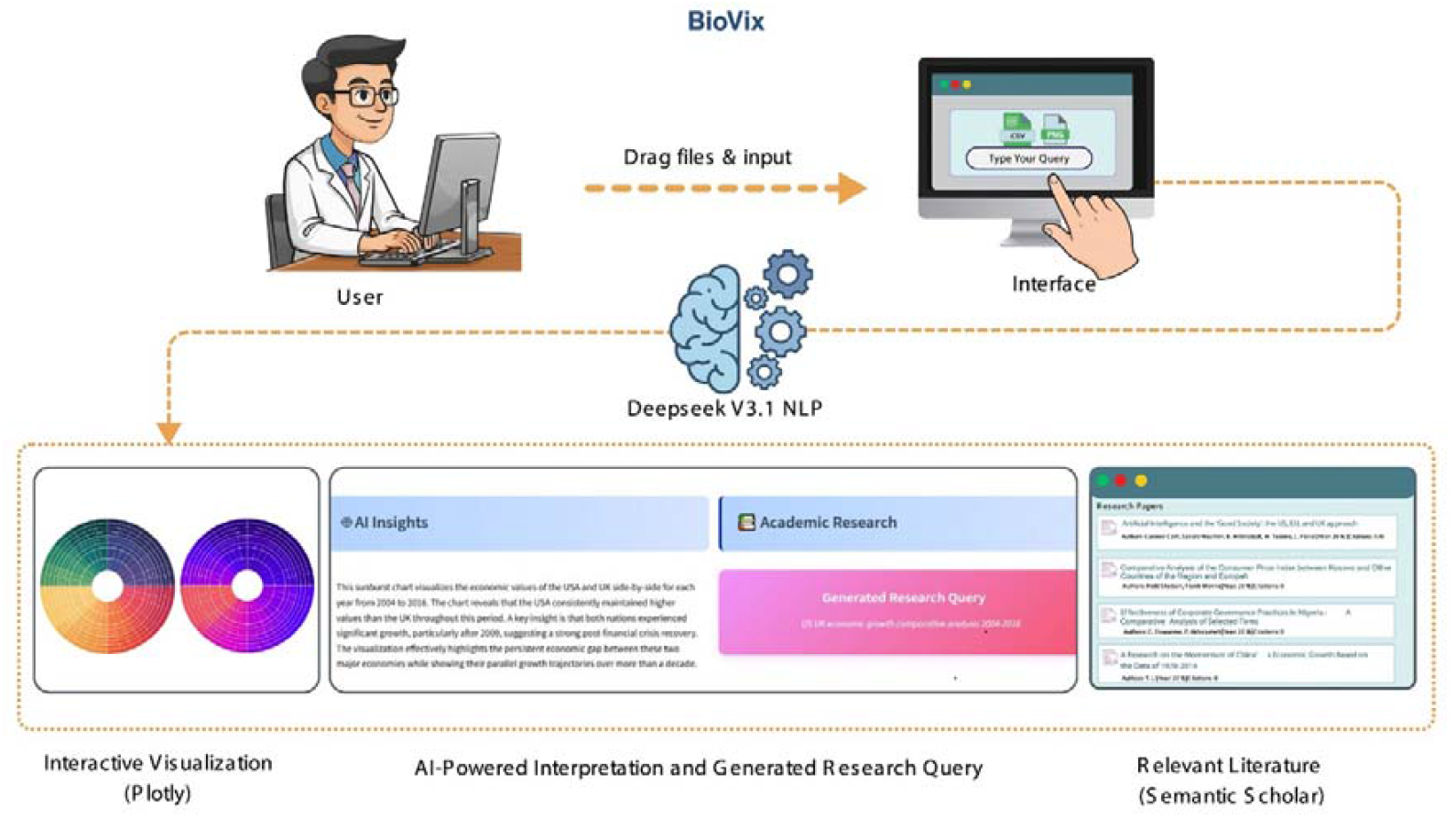
User interaction flow in BioVix, from data or chart upload and natural-language queries to visualization, AI-driven interpretation, and literature support.

## 2. Materials and Methods

### 2.1. Architecture

The BioVix architecture consists of three functional modules: (i) a visualization-to-literature-search workflow, (ii) an interpretation of uploaded graphs with user queries, and (iii) conversational support for uploaded data. The architectural structure of BioVix is further described in detail below.

#### 2.1.1. Visualization-to-literature-search workflow

The visualization-to-literature-search workflow comprises several steps **(Figure 2)**, including code generation and correction, generated chart interpretation, search query formulation, and Semantic Scholar integration.

**Figure 2.**
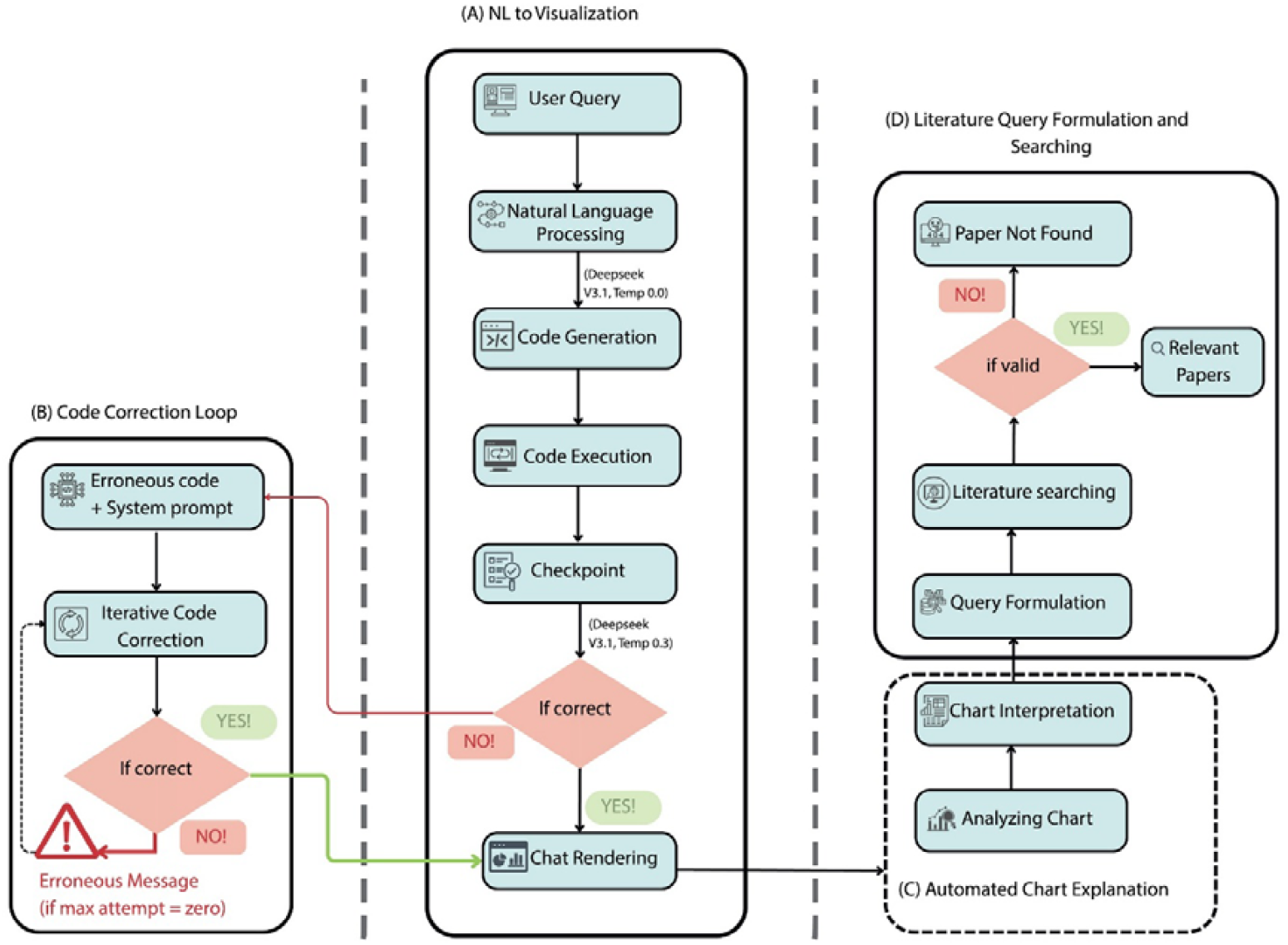
Schematic representation of the BioVix visualization framework supporting literature-driven validation of results.

##### i. Code generation and correction

A function named “*generate_chart_code”* was developed to process NLQs into an executable Python script for chart generation. BioVix architecture processes the input by concatenating the user’s NLQ with an engineered prompt. This prompt includes a system role defining data visualization assistant capabilities and constraints, metadata extracted from the uploaded dataset (such as header names and column data types), the user’s NLQ, and an output specification that enforces the use of the Plotly package for graph creation. This code-generation function invokes DeepSeek V3.1 LLM to generate Python code using a zero-temperature setting. The “*fix_chart_code*()” function was developed to improve inconsistent and erroneous code for the visualization by employing an iterative error-correction technique [17]. The generated Python code is executed within an isolated environment for safety. Upon execution failure, BioVix enters a debugging loop with up to 3 iterations. During this loop, the function engineered a refined correction prompt using three key inputs: (i) the original failed code, (ii) the exception message received during execution, and (iii) metadata describing the uploaded data elements. The prompt incorporates logical constraints to ensure that the DeepSeek V3.1 LLM does not fabricate data and verifies that all required data columns are adequately loaded before attempting code correction. The model then iteratively applies this prompt up to a three-iteration limit to resolve the error. The code-correction function follows the same protocol as the code-generation process, using a zero-temperature setting for the DeepSeek V3.1 LLM [20].

##### ii. Generated chart interpretation

After effective visualization, BioVix generates NL explanations detailing the content and trends of generated charts. The function “*generate_explanation()”* converts the visualization into a logical NL summary that expresses key insights of the generated chart. This function takes several inputs, including: (i) user query, (ii) the generated code, (iii) metadata, and (iv) statistical summaries of the uploaded data. This information is used to engineer a prompt that guides the DeepSeek V3.1 LLM in processing the data, identifying essential trends and patterns, and providing a concise interpretation [22].

##### iii. Search query formulation and semantic scholar integration

An automated literature search to validate visualization results against existing academic articles is a distinctive feature of BioVix. The function “*analyze_explanation_for_query()”* was developed to convert a visualization explanation into a relevant academic search query tailored to the specific context. This function requires three primary inputs: (i) the user query, (ii) metadata from the uploaded data, and (ii) the chart explanation rendered in the preceding module. An engineered prompt is used to instruct DeepSeek V3.1 LLM to produce a specific query that integrates methodological terms and uploaded data-related elements. This step uses a low temperature of 0.1 to mitigate randomness and ensure a precise output. The resulting formulated query enables BioVix to associate observed visualization trends with relevant existing literature [12]. The function “search_semantic_scholar()” was developed to send the formulated query to the Semantic Scholar API. BioVix retrieves the five most relevant and up-to-date articles from the academic literature repository using the Semantic Scholar database. To ensure reliable retrieval performance and minimize network or server-related issues, the BioVix framework incorporates features such as exponential back-off for connection aborts and automatic control of connection rate limits. (**Table 1)** contains detailed features for literature articles retrieval.

**Table 1.** Parameters defined in BioVix for the literature search process.

| PARAMETERS | VALUE | REASON |
| --- | --- | --- |
| Results Limit | Five | To prioritize the most relevant |
| Sorting | Relevance | To validate user data with the literature |
| Retrieved information | Title, authors, year, abstract, citation no, URL | Metadata for preview |
| Connection time | Exponential pull back | Handling rate limits |
| Connection retries | Three | Balance between persistence and wait time |

#### 2.1.2. Multimodal chart interpretation

BioVix provides an AI-based chart interpretation function. The “*interpret_graph_with_qwen()”* function was developed to allow users to upload a chart in PNG, JPG, and JPEG formats for AI-powered interpretation. It takes two inputs: (i) the binary of the uploaded image to ensure compatibility with the large vision-language model, and (ii) an NLQ by the user. A crafted prompt then combines textual and visual components to instruct the Qwen2.5-VL-32B-Instruct LLM to identify insightful patterns and outliers in the provided chart [24]. The Qwen2.5-VL-32B-Instruct LLM was implemented with a temperature setting of 0.3, ensuring consistent and contextually grounded outputs [25].

#### 2.1.3. Conversational feature

BioVix provides a conversational feature for uploaded datasets that enables interactive and intelligent reasoning by allowing users to identify patterns in the data using NLQs. This module takes two inputs: (i) the user query and (ii) a well-structured dataset summary that includes the number of rows and columns, file headers, data samples (first five rows), statistics, and missing values. An engineered prompt then guides the GPT-OSS-20B LLM to identify patterns, trends, and key insights in the data. The GPT-OSS-20B LLM is invoked with a temperature setting of 0.7 to promote individualized and inquisitive responses suitable for conversational interaction. This NL-based description helps bridge the gap between raw data findings and AI-driven interpretation.

### 2.2. Web-based graphical user interface

The web-based graphical user interface (GUI) of BioVix was implemented using the Python package Streamlit [26]. The web GUI layout features a sidebar panel that allows users to upload the sample data for quick evaluation. The GUI was further designed with two expanders, which sequentially house the conversational feature and the multimodal chart interpretation module. The Plotly package was used as the primary visualization library to create diverse and interactive visualizations with complex NLQs.

### 2.3. Large Language Models (LLMs) selection

BioVix employs different LLMs for distinct functions instead of relying on a single LLM for all tasks. We selected the task-specific LLM model selection strategy to achieve higher accuracy, efficiency, and interpretability, while reducing the risk of hallucinations and excessive token usage [27]. This strategy reflects the fact that different functions of BioVix architecture, such as visualization-to-academic search integration, conversational feature, and multimodal processing, require diverse linguistic and reasoning capabilities. The corresponding characteristics of selected LLMs and their functions in BioVix are detailed in **Table 2**.

**Table 2.**
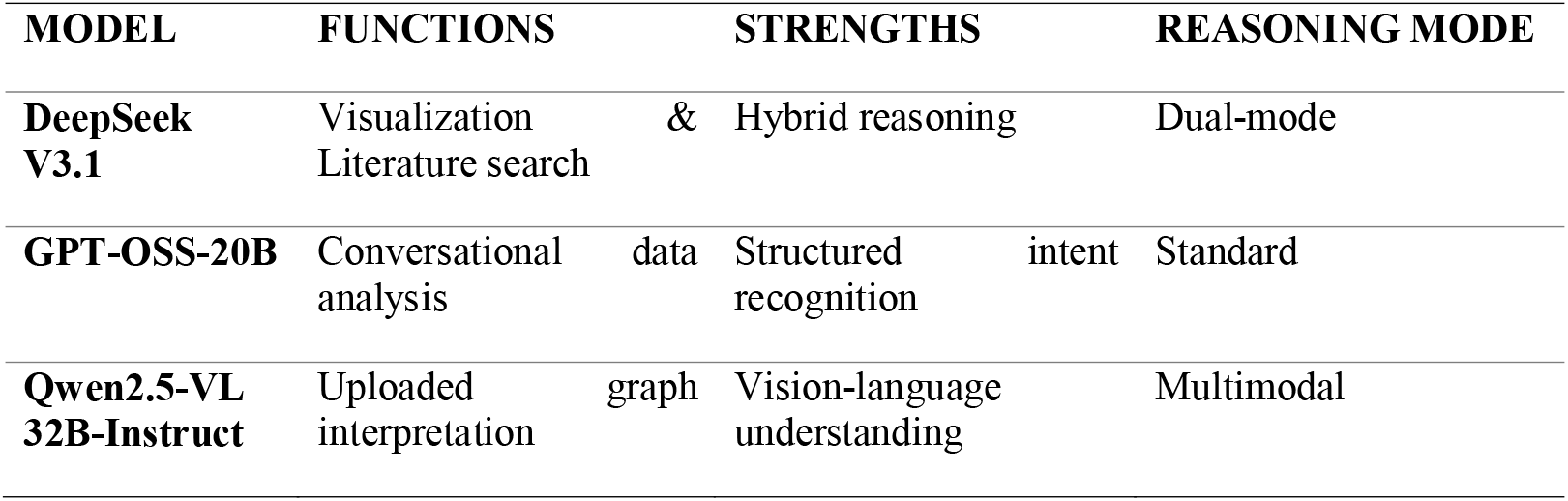
Qualitative assessment of the three large language models selected for implementation in BioVix.

### 2.4. Datasets used for evaluation

We downloaded three different datasets to evaluate the efficiency and accuracy of BioVix. These datasets include the protein expression data from Human Proteome Map (HPM) from GeneRanger (https://generanger.maayanlab.cloud/download), genomic peak annotation data from Human Peripheral Blood Mononuclear Cells (PBMCs) generated through ATAC sequence analysis from 10xGenomics (https://www.10xgenomics.com/datasets/10-k-human-pbm-cs-atac-v-1-1-chromium-x-1-1-standard-2-0-0), and a clinical diabetes dataset from the Pima Indians Diabetes Database available on the open source platform Kaggle (https://www.kaggle.com/datasets/uciml/pima-indians-diabetes-database/data). We accessed these datasets on 03-01-2026. A brief description of each dataset is provided below:

#### 2.4.1. Pima Indians diabetes database

The Pima Indians diabetes database is a publicly available dataset that consists of diagnostic variable values from adult female patients of Pima Indian heritage, for predicting the symptoms of diabetes in patients [28]. The dataset contains a total of 768 records with 9 attributes, such as pregnancies, glucose concentration, blood pressure, skin thickness, insulin levels, body mass index, and age. A binary classification variable indicates the diabetes status in females. Due to the well-structured and well-annotated labels, this dataset is often used for medical data analysis and benchmarking [28].

#### 2.4.2. Human peripheral blood mononuclear cells (PBMCs)

The 10k Human PBMCs ATAC v1.1 dataset contains data from approximately 10,000 human PBMCs. This is publicly available single-cell epigenomics dataset that was generated using the 10x Genomics Chromium X platform [29]. The data was processed using Cell Ranger ATAC version 2.0.0, and the peak annotation file (CSV format) was used in this study. The peak annotation file contains a total of 179,416 rows and 6 columns, such as chromosome number (chrom), genomic coordinates of accessible regions (start and end), along with respective gene annotations, including gene id, distance from TSS (Transcription Starting Site), and peak type to facilitate epigenetic profiling.

#### 2.4.3. Human proteome map

Proteomics datasets processed by GeneRanger were retrieved from the Human Proteome Map database, which provides large-scale protein expression abundance profiles across 23 human tissues and 6 cell types with 17,294 records [30]. These datasets consist of processed protein-level expression data, allowing direct downstream analysis without raw mass spectrometry preprocessing.

## 3. Results

### 3.1. Overview of BioVix

BioVix is an innovative web-based application that aims to simplify data visualization by integrating LLM-driven visualization with academic literature validation. It combines various functions into a single, user-friendly framework, thereby eliminating the need for extensive technical expertise for data visualization (**Figure 3**). BioVix utilizes Streamlit for the web-based user interface and Plotly package as the primary visualization library. The OpenRouter API (Application Programming Interface) is used to access the specified LLMs (DeepSeek V3.1, Qwen2.5-VL-32B-Instruct, and GPT-OSS-20B), which are used, respectively, for the visualization-to-literature-search workflow, an interpretation of uploaded graph based on user query, and conversational support for uploaded data. BioVix uses the Semantic Scholar API to retrieve the most relevant literature. A key feature of BioVix is automated visualization code generation, which allows users to create interactive charts without writing any code. The system automatically generates and integrates the required code modules to produce figures. Moreover, BioVix uses robust AI-based chart-interpretation systems, that enable users to obtain contextual insights from visualizations. Another novel feature of BioVix is the integration of academic literature search into the visualization process. BioVix integrates with platforms such as Semantic Scholar to provide real-time, evidence-based validation by cross-referencing critical data insights with existing published articles. This unique step ensures that users, especially researchers and domain experts, can summarize and support their findings with legitimate current academic evidence.

**Figure 3.**
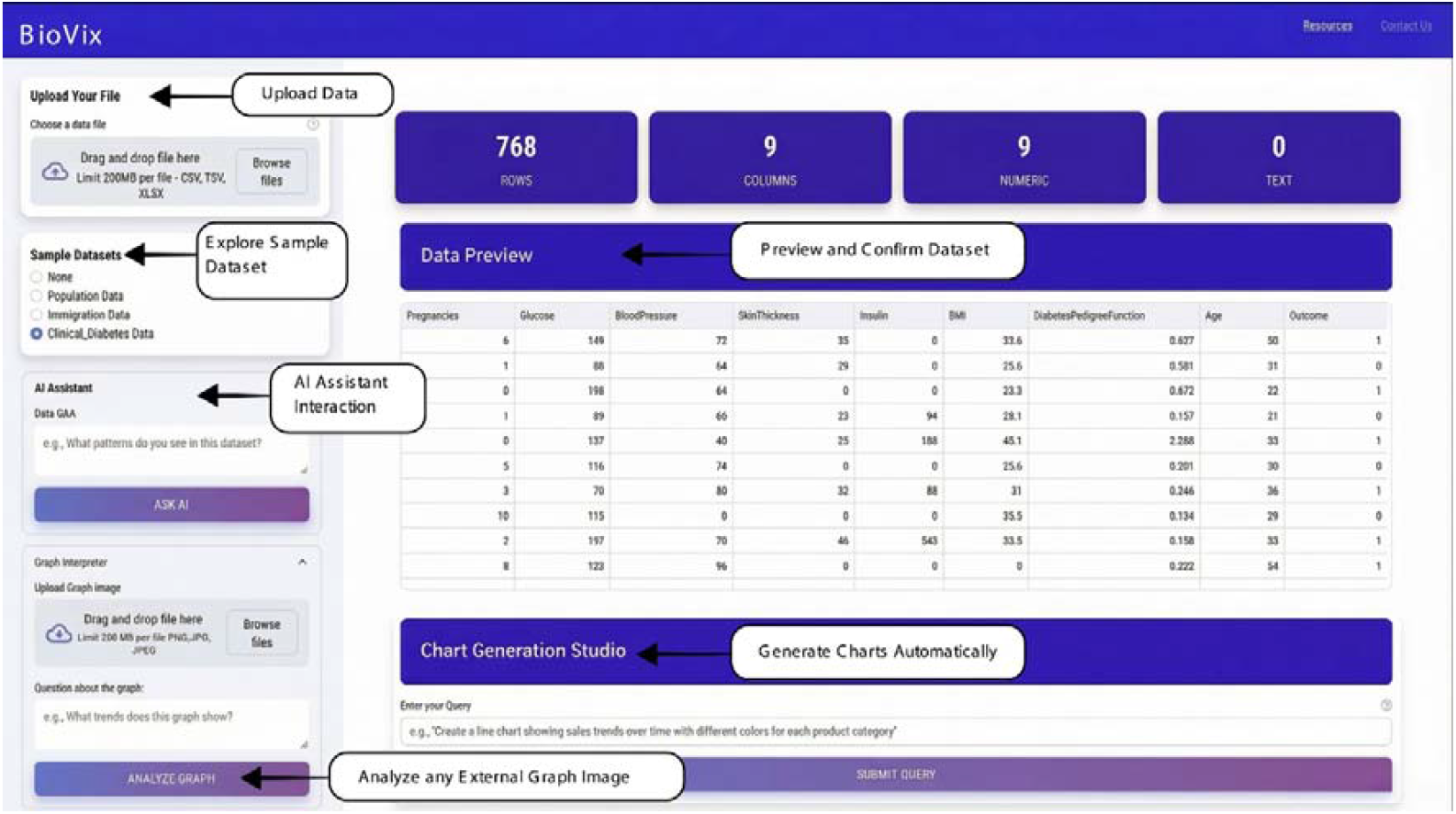
BioVix web graphical user interface demonstrating the integrated workflow from data upload and dataset validation to AI-assisted analysis, visualization generation, and external graph interpretation.

### 3.2. Evaluation of BioVix

We evaluated the effectiveness and reliability of BioVix outcomes using diverse datasets. The evaluation focused on three core aspects of the BioVix workflow: code generation success, interpretive accuracy, and literature retrieval relevancy. Across evaluations, BioVix consistently demonstrated robust alignment between visualization-derived trends, AI-generated textual interpretations, and findings reported in current scholarly literature. Collectively, these three case studies demonstrate that BioVix generalizes effectively across heterogeneous biological datasets, including proteomics, epigenomics, and clinical data, while maintaining consistent visualization quality, interpretability, and literature validation.

#### 3.2.1. Case study 1

The first evaluation was conducted by using the Human Proteome Expression dataset obtained from GeneRanger. The results produced by BioVix when using this data are presented in **Figure 4**. A representative subset of the uploaded dataset is shown in **Figure 4A**, which includes the protein identifiers (genes) along with their quantified expression values across various tissues and cell types at the gene level. A hierarchical sunburst graph **(Figure 4B)**, illustrating the distribution of protein expression abundance across grouped tissue and cell categories. This visualization was generated in response to the natural-language query (NLQ): “*Create a Plotly sunburst using only tissue group to tissue hierarchy with summed expression*”. The chart enables investigation of protein expression distribution in adult, fetal, immune tissues, and further biological profiling. This visualization demonstrates BioVix’s ability to transform complex, multidimensional datasets into clear and interpretable hierarchical representations. Gene-level protein expression across a wide range of tissues and cell types is further illustrated using a stacked bar plot **(Figure 4C)**. This representation allows the comparison of cross-tissues and cross-cells and has shown variability in protein abundance patterns, where some genes show broad expression while others exhibit tissue-enriched patterns. This plot was generated using the NLQ: “*Create the stacked bar plot for top 10 highly expressed genes across different tissues and cells with distinct color coding for different genes*”. The top expressed genes in terms of average abundance are shown in **Figure 4D** with a bar chart and color encoded to highlight the magnitude of relative expression based on the NLQ: “*Plot a bar chart of the top 10 genes ranked by average expression with different dark colors for values*”. This visualization helps quickly identify highly abundant proteins and strengthens the ability of BioVix to aggregate and rank data according to exploratory queries. **Figure 4E** presents the automated insight and semantic research query out of the visualized data shown in **Figure 4C**. BioVix produced an objective summary of observed protein expression trends across human tissues and formulated a research query reflecting tissue-specific expression patterns suitable for downstream literature exploration. The retrieved scholarly articles are displayed in **Figure 4F**. These articles align with the main themes of the visualization, such as the regulation of tissue-specific genes and protein expression. This correspondence demonstrates that BioVix can be useful in the process of transforming the qualitative or quantitative characteristics of visualizations into semantically meaningful literature suggestions, and thus has the capacity to mediate the gap between exploratory data analysis and specific domain knowledge discovery.

**Figure 4.**
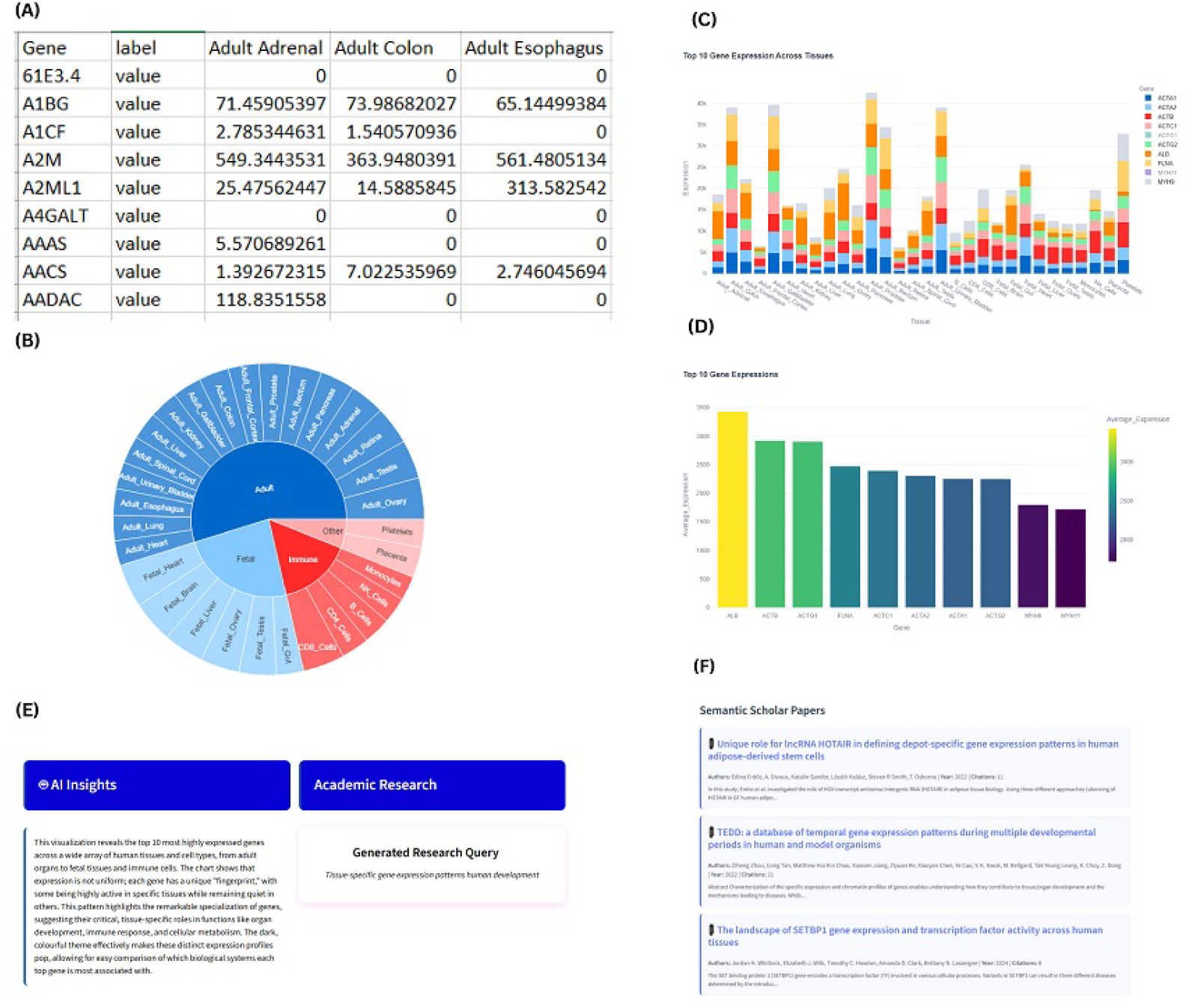
BioVix results derived from the Human Proteome Map (HPM) protein expression dataset in response to user-defined natural-language queries. (A) Representative subset of the uploaded dataset. (B) Sunburst visualization illustrates hierarchical protein expression across tissue groups. (C) Stacked bar plot showing protein expression levels across tissues and cell types. (D) Bar chart of the ten most highly expressed proteins at the gene level. (E) AI-generated insights derived from the visualization and the automatically formulated research query. (F) Relevant scholarly literature retrieved via Semantic Scholar in response to the generated research query.

#### 3.2.2. Case Study 2

In this evaluation, the performance of BioVix was assessed using a genomic peak annotation dataset derived from 10x Genomics ATAC-seq data of 10,000 human peripheral blood mononuclear cells (PBMCs). The results of this analysis are presented in **Figure 5**. The ATAC processed dataset uploaded is presented in **Figure 5A**, as a genomic peaks annotation file including genomic coordinates, associated genes, distance to transcription start sites, and peak classification (distal, promoter, or intergenic). Summary bar plots illustrating peak distributions are shown in **Figure 5B**. These plots were generated in response to the natural-language query (NLQ): “*Two side-by-side bar charts on white: Peak Counts by Chromosome (descending, multicolored bars) and Peak Counts by Peak Type (blue bars: distal highest, promoter smaller, intergenic minimal)*”. The chromosome-wide distribution showed that there are variations in the density of regulatory peaks, while the distribution by peak type shows a substantially higher number of distal regulatory elements compared with promoter and intergenic peaks. These visual summaries demonstrate the ability of BioVix to quickly evaluate the global landscape of uploaded datasets. A Funnel graph **(Figure 5C)** that shows the relationship between chromosomes and types of peaks generated generated using the NLQ: “*Create a Plotly funnel chart where peaks from chromosomes chr1 flow through stages distal, promoter, and intergenic, each chromosome shown as a colored segment with counts and connecting flows between stages, final counts at intergenic, and a legend mapping chromosomes to colors*”. This visualization highlights the distribution of the various peak types across chromosomes, emphasizing the prevalence of distal peaks and their broad genome-wide coverage. A hierarchical tree-map graph **(Figure 5D)** with the relationship between chromosomes and peak types was generated using the NLQ : “*Create treemap for different peaks (promoter, distal and intergenic) distribution over the different chromosomes with different colors*”. The tree-map also allows comparative exploration of regulatory element abundance across chromosomes, while retaining similar categorization within each chromosomal group. The automatically generated AI insight summary and semantic research query derived from the visualization shown in **Figure 5B** are presented in **Figure 5E**. These insights provide a high-level understanding of chromosomal distribution and regulatory peak classifications without reliance on gene-specific terminology. The research question that is associated with the observed patterns generalizes the trends in a field-related formulation that aims at the distribution of chromosomes and classification of regulatory peaks. **Figure 5F** represents the research articles retrieved via Semantic Scholar based on the generated query. The retrieved literature aligns closely with the visualization themes, including promoter-enhancer interactions, chromatin organization, and regulatory variant mechanisms. This demonstrates that BioVix can translate visualization features into semantically meaningful literature recommendations, which facilitates integrative analysis.

**Figure 5.**
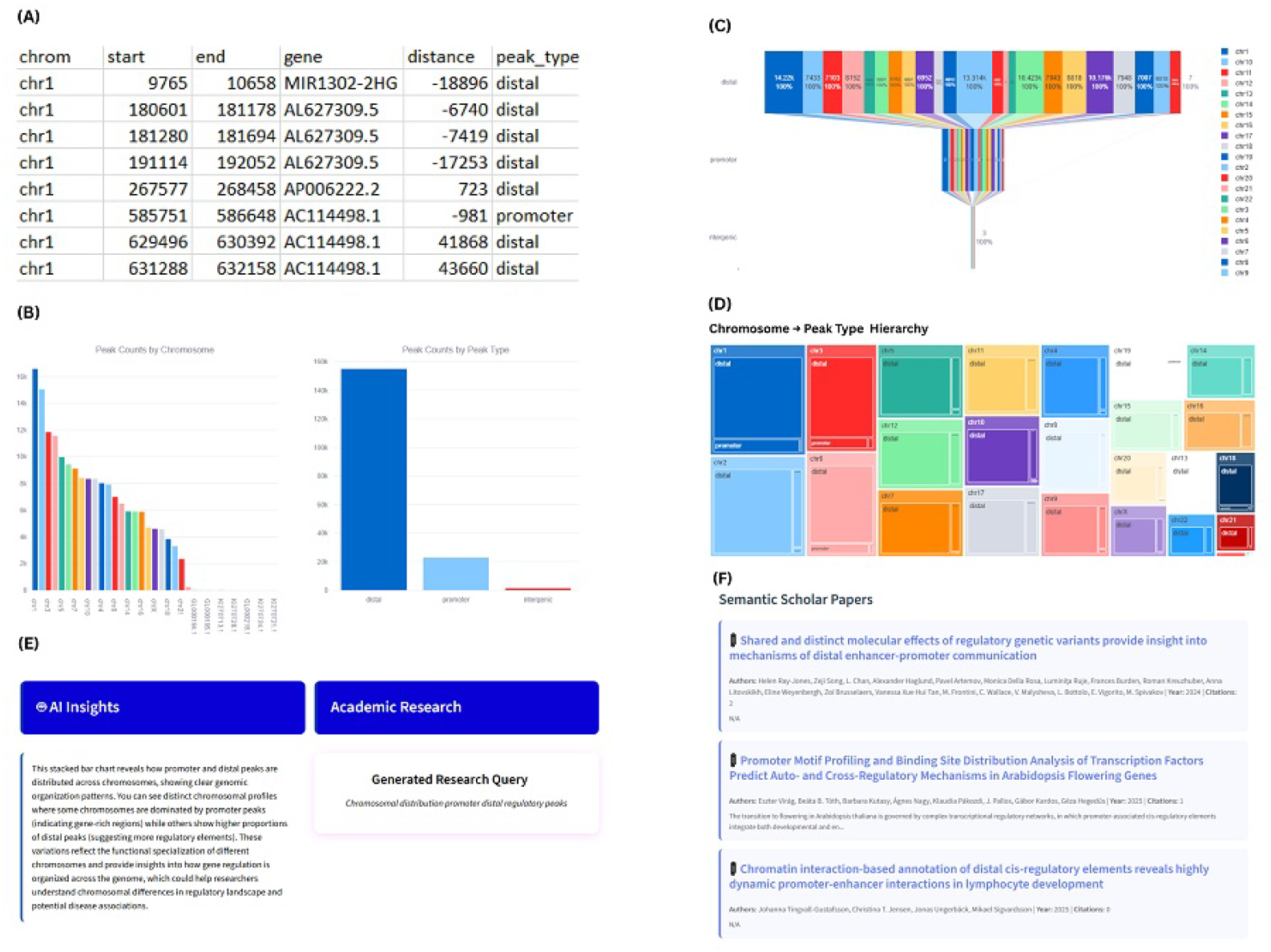
BioVix results for the genomic peak annotation dataset generated in response to user-submitted natural-language queries. (A) Representative subset of the uploaded peak annotation dataset. (B) Bar plots showing the distribution of peak counts by chromosome and by regulatory peak type. (C) A funnel diagram illustrates the occurrence and flow of peaks across chromosomes by peak category. (D) Hierarchical treemap depicting the relationship between chromosomes and regulatory peak types. (E) AI-generated insights derived from the visual analyses. (F) Relevant scholarly literature retrieved via Semantic Scholar based on the automatically generated research query.

#### 3.2.3. Case Study 3

In this case, we used the clinical diabetes dataset, which contains clinical data of female patients aged 21 or older of Pima Indian heritage. The results generated by BioVix using this dataset are shown in **Figure 6. Figure 6A** displays a representative subset of the input dataset, which consists of physiological and clinical predictor variables, including the number of pregnancies, plasma glucose concentration, blood pressure, skin thickness, serum insulin, body mass index (BMI), diabetes pedigree function, and age. The dataset includes a binary outcome variable indicating diabetes diagnosis (0 = non-diabetic, 1 = diabetic). To explore multivariate relationships, the following natural-language query (NLQ) was submitted to BioVix: “*Build a 3D scatter plot using Glucose, BMI, and Age as axes, and color the points by diabetes Outcome*”. This request represents a high-complexity task due to the exigencies of 3D rendering, attribute mapping, and categorical coloring. **Figure 6B** presents a BioVix-generated 3D scatter plot that accurately positions data points along the BMI, blood pressure, and age axes, uses a binary color code to distinguish the outcomes, and presents the indicative health risk of elevated BMI and blood pressure correlated with diabetic outcomes in old age. This multivariate visualization emphasizes BioVix efficiency in dimensionality reduction and in visualizing multivariate correlations for clinical datasets. **Figure 6C** presents a donut chart that contains the proportion of individuals, clearly differentiates both categories with different colors. BioVix highlights the imbalance in understanding dataset composition before model training or association analysis. **Figure 6D** display the scatter-matrix plot that describes pairwise relationships among all clinical variables. Each subplot displays the distribution of two predictors, with density contours indicating linear or non-linear relationships. The matrix reveals heterogeneous relationships across features, with some variable pairs showing weak correlations and others demonstrating stronger associations, particularly involving glucose, BMI, insulin, and age. **Figure 6E** illustrates AI-assisted explanations and corresponding research queries for the **Figure 6C** visualization, emphasizing the prevalence of diabetes in the dataset and identifying variables that may be associated with it (glucose, insulin, BMI, age). BioVix formulates a specific research query as: “Diabetes prevalence risk factors pregnancy glucose insulin BMI “, which is then used to search the related literature. Finally, **Figure 6F** displays the resulting literature recommendations retrieved via Semantic Scholar for the above-formulated query **Figure 6E**. The retrieved articles address gestational diabetes, postpartum glucose tolerance, risk factors for microalbuminuria, and related metabolic disorders. These articles align closely with the AI-formulated query and the observed visualization patterns, demonstrating BioVix’s capability to derive coherent scientific context from data visualizations and to bridge exploratory analysis with relevant domain literature.

**Figure 6.**
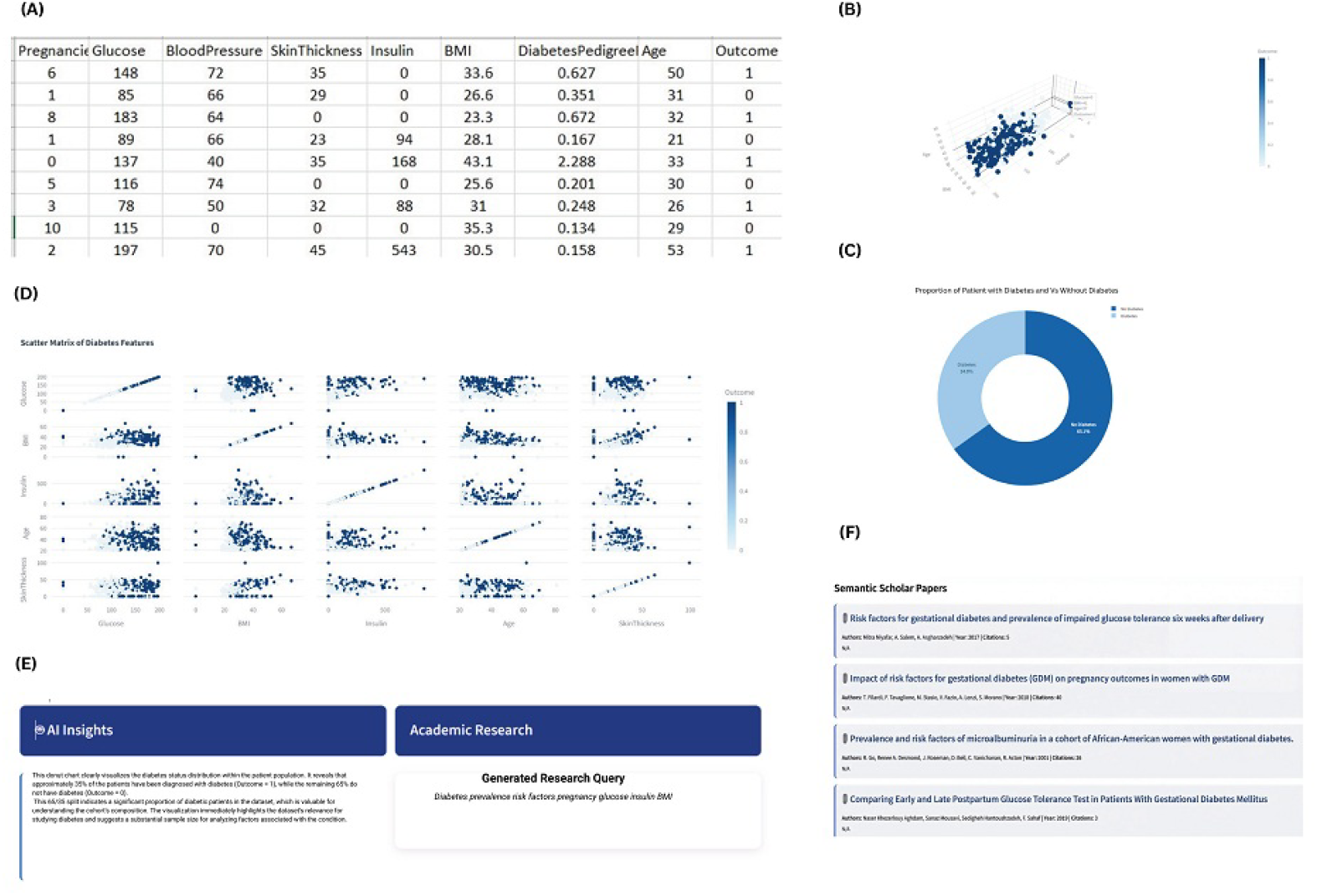
Exploratory analysis of diabetes-related clinical data using BioVix. (A) Summary of patient clinical features in the diabetes dataset. (B) Three-dimensional scatter plot illustrating multivariate relationships among clinical features. (C) Distribution of diabetes outcome classes. (D) Pairwise scatter plots depicting correlations among clinical variables. (E) AI-generated insights and automatic research query formulation based on the visual analyses. (F) Relevant scholarly literature retrieved via Semantic Scholar in response to the generated research query.

## 4. Conclusion

In this study, we developed BioVix, an LLM-based visualization tool designed to bridge the gap between exploratory data visualization and scholarly validation. BioVix adopts a multi-model architecture that overcomes the limitations of single-model systems by leveraging DeepSeek V3.1 for code and logic generation, Qwen2.5-VL for multimodal interpretation, and GPT-OSS-20B for conversational analysis. We evaluated BioVix on three diverse biological datasets: human proteome abundance profiling, epigenomic peak annotation (ATAC-seq), and clinical diabetes diagnosis. The results indicate that BioVix effectively handles heterogeneous data types, producing complex visualizations while supporting contemporary literature-based validation. These evaluations further demonstrate BioVix’s capability to support diverse analytical tasks across protein expression, peak annotation, and clinical diabetes datasets. This platform uniquely provides an automated pipeline for searching relevant academic literature to corroborate research results. BioVix enhances data interpretation by integrating automated visualization, analytical reasoning, and literature search within a unified workflow. It is designed to favor two-way analysis, allowing users to transition freely between raw data and visual displays, and between visualizations and actionable insights. While the platform substantially reduces programming effort, effective use still requires domain expertise, informed selection of visualization strategies, and careful interpretation of results. Importantly, as BioVix is an LLM-based system, all generated visualizations, interpretations, and literature recommendations should be independently verified by users prior to incorporation into scientific conclusions or publications. This ensures responsible use and mitigates the risk of inaccuracies inherent to generative models. Future improvements should prioritize expanding the range of supported visualization libraries by incorporating packages such as Matplotlib and Seaborn, and by using appropriate dependency management to support a broader range of chart types. Additional improvements include automatically refining generated graphs in response to follow-up user queries for dynamic exploration, and targeting enhancements at preprocessing deposited raw datasets and granting access to a greater number of academic repositories to enable more comprehensive literature validation. The integration of these capabilities positions BioVix as a promising platform for data-driven research workflows.

## Acknowledgment

Not applicable.

## Author contributions

M.Z.B. led the development of the BioVix framework, including software implementation, system integration, testing, and manuscript preparation. R.S.A. contributed to logical programming, software development, visualization, testing, and manuscript writing. E.F. performed validation, benchmarking, figure generation, and manuscript editing. M.T.Q. conceptualized and supervised the study, provided resources, contributed to logical programming, and critically revised the manuscript.

## Data Availability

No new data were generated in this study. All datasets used for evaluation are publicly available, including Human Proteome Map protein expression data from GeneRanger, ATAC-seq PBMC peak annotation data from 10x Genomics, and the Pima Indians Diabetes Database from Kaggle. BioVix is accessible via Hugging Face, and the full source code and installation instructions are available on GitHub.

## Ethical Statement

This study did not involve any experiments with animal subjects or human participants. The authors utilized ChatGPT v2 to enhance the clarity and readability of the code and to polish the language of the manuscript. All content generated through this service was carefully reviewed and edited by the authors, who take full responsibility for the final content of the publication.

